# Energetical equivalence between air resistance and gradients in running

**DOI:** 10.1101/2023.06.01.543316

**Authors:** M. Leclerc

## Abstract

For a given running speed and for any wind speed, we calculate the slope for which the effort to overcome in the absence of wind is energetically equivalent to running with the original wind speed on a flat track. The influence of headwind and tailwind is thus made numerically comparable to the influence of a positive or negative slope. The same applies to the lack of air resistance on a treadmill and its compensation by adjusting the incline. Moreover, for turning point routes physiological corrections are considered and the impact of speed adjustments is analyzed.

## 1 Calculation of air resistance and gravitational force

The force induced by air resistance can be described in the following way:

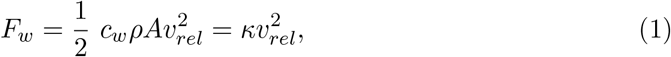

where *ρ* denotes the density of the air, *A* the cross section of the runner perpendicular to his motion, *c*_*w*_ the so called drag coefficient and *v*_*rel*_ the relative velocity between the runner and the air. Our considerations will be limited to wind directions parallel to the running direction, i.e., tailwind or headwind. For any other wind direction, the results are applicable to the component of the wind speed which is parallel to the running direction.

Under usual climatic conditions, we have *ρ* = 1.2 kg*/*m^3^. For the cross section, i.e., the projected area perpendicular to the direction of the wind speed and of the runner’s motion, values between 0.5 m^2^ and 0.7 m^2^ can be assumed, while the drag coefficient *c*_*w*_ has been determined to 0.8 − 0.9, see, e.g., [1] or [2]. The cross section depends on the stature and posture, the drag coefficient on clothing. In this article, we will use the values *A* = 0.5 m^2^ and *c*_*w*_ = 0.9. The constant *κ* introduced in Eq. (1) then takes the value of about 0.27 kg*/*m.

The component tangential to the motion of the runner induced by the gravitational field while running on a gradient is given by:

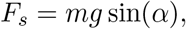

where the angle of slope *α* is defined as 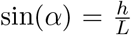, with the covered height difference *h*, the covered distance *L*, the runner’s mass *m* and the the strength of the gravitational field *g*. For small angles we have approximately sin(*α*) ≈ *α* ≈ tan(*α*). On the other hand, the slope (in percent) is defined by *s* = 100 tan(*α*). We can thus write:

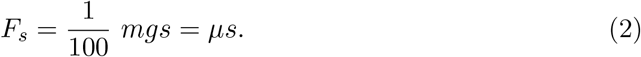

The approximation sin(*α*) ≈ *α* ≈ tan(*α*) is valid with sufficiently high accuracy for slopes up to 25 − 30 % and will be used throughout this article. In the numeric examples, we further use the value *m* = 70 kg. With *g* = 9.81 m*/*s^2^, this results in *µ* = 6.87 N for the constant introduced in Eq. (2).

The resistive force corresponds the energy input per meter, or more precisely, in the cases considered here, to the additional energetical effort (per meter) compared to a flat and windless track. The additional power, i.e., the additional energetical effort per second, is, in addition, proportional to the running speed. The comparison between wind and gradient made in this article always refers to a constant speed *v*, which means that our considerations can be made either based on force or on power.

In our calculations of equivalent slopes in the following sections, the constant *κ/µ* plays a central role. It has a dimension of *s*^2^*/m*^2^ and reads explicitly:

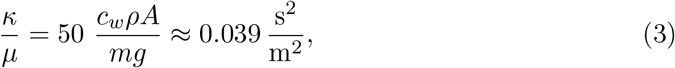

where the numeric value corresponds to the assumptions *m* = 70 kg, *A* = 0.5 m^2^ and *c*_*w*_ = 0.9. Quite generally, all the results of the following sections can be directly rescaled to different values for *m, A* and *c*_*w*_ by considering the respective proportionality.

The concept of equivalent slopes in describing the influence of air resistance on a runner has been introduced by Hill, see [1]. In this article, we apply this concept to several specific situations and provide results in form of tables and plots, suitable for direct reference.

## 2 Equivalent slope on a treadmill

In this section, we determine the incline that has to be set on a treadmill in order to compensate for the lack of air resistance. We calculate the slope for which the tangential component of gravity, Eq. (2), is equal to the force of air resistance, Eq. (1), that would have to be faced by a runner at a given speed while running outdoors on a flat and windless track, i.e., *µs* = *κv*^2^ and thus

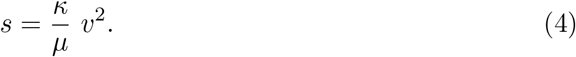

Using the values given in section 1 for *κ* and *µ*, we find the slope percentages displayed in Table 1. For reference purposes, we also indicate the corresponding angles of slope (in degrees). Similar calculations have been performed in [1], but based on a cross section of about 0.6 m^2^, leading to slightly higher values. In any case, the popular suggestions to set the incline to 1 − 2 % seem to be slightly exaggerated for most runners.^1^

**Table 1:**
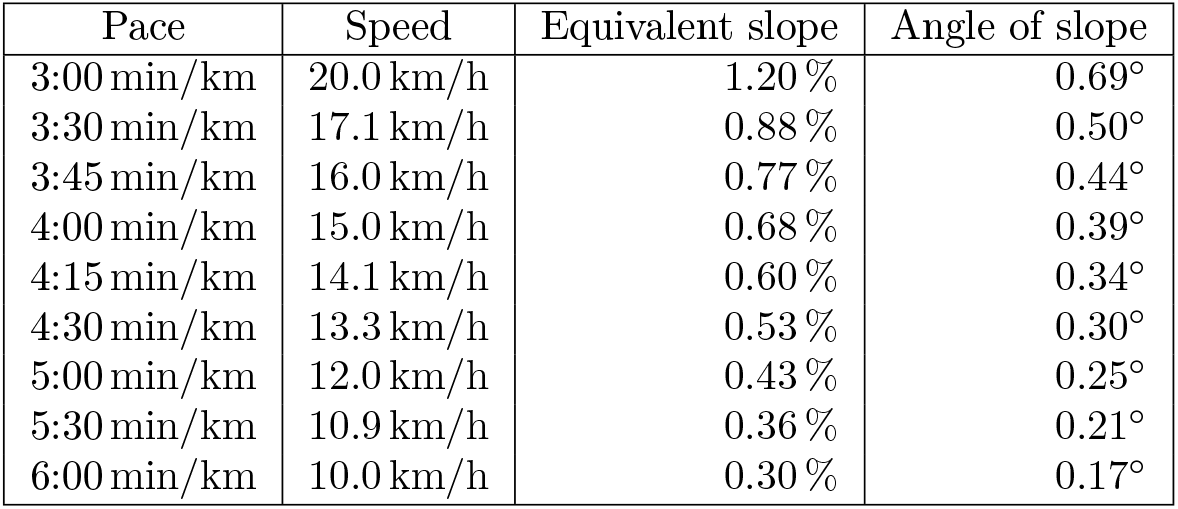
Equivalent slope on a treadmill

| Pace | Speed | Equivalent slope | Angle of slope |
| --- | --- | --- | --- |
| 3:00 min/km | 20.0 km/h | 1.20 % | 0.69° |
| 3:30 min/km | 17.1 km/h | 0.88 % | 0.50° |
| 3:45 min/km | 16.0 km/h | 0.77 % | 0.44° |
| 4:00 min/km | 15.0 km/h | 0.68 % | 0.39° |
| 4:15 min/km | 14.1 km/h | 0.60 % | 0.34° |
| 4:30 min/km | 13.3 km/h | 0.53 % | 0.30° |
| 5:00 min/km | 12.0 km/h | 0.43 % | 0.25° |
| 5:30 min/km | 10.9 km/h | 0.36 % | 0.21° |
| 6:00 min/km | 10.0 km/h | 0.30 % | 0.17° |

## 3 Equivalent slope in headwind and tailwind

We now consider a runner moving with velocity *v* on a flat track. Let *w* denote the wind speed, positive values representing tailwind and negative ones representing headwind.^2^ The relative velocity between the runner and the air environment is given by *v*_*rel*_ = *v* −*w*.

The air resistance for this case reads

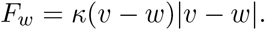

This results from Eq. (1) taking into account the fact that, whenever *w* exceeds *v*, the force points in the direction of the runner and thus changes sign. In the absence of wind, the air resistance is given by

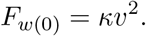

Our aim is to derive the slope *s* resulting, in the absence of wind (*w* = 0), in the same energetical expenditure than running with wind (*w* ≠ 0) on a flat track, i.e., the slope for which the gravitational force tangential to the running direction is equal to the additional force *F* = *F*_*w*_ − *F*_*w*(0)_.

Equating the additional force to *F*_*s*_ from Eq. (2), we find the equivalent slope in the form:

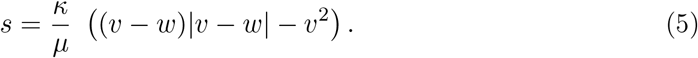

For *w <* 0, we have *s >* 0 (headwind), while for *w >* 0 (tailwind), we have *s <* 0, this being independent on whether *w > v* or not. This simply reflects the fact that tailwind results in less effort compared to the standard (flat and windless) case.

Table 2 and the chart in Fig. 1 show the results for runner running at a pace of 3 min*/*km, 4 min*/*km, 5 min*/*km or 6 min*/*km, i.e., at a speed of 20 km*/*h, 15 km*/*h, 12 km*/*h or 10 km*/*h, using the values for *κ* and *µ* given in section 1.

**Table 2:** Equivalent slope *s* in headwind and tailwind

| $w$ | 3:00 min/km | 4:00 min/km | 5:00 min/km | 6:00 min/km |
| --- | --- | --- | --- | --- |
| −40 km/h | 9.63 % | 8.43 % | 7.70 % | 7.22 % |
| −30 km/h | 6.32 % | 5.42 % | 4.88 % | 4.51 % |
| −20 km/h | 3.61 % | 3.01 % | 2.65 % | 2.41 % |
| −15 km/h | 2.48 % | 2.03 % | 1.76 % | 1.58 % |
| −10 km/h | 1.50 % | 1.20 % | 1.02 % | 0.90 % |
| −5 km/h | 0.68 % | 0.53 % | 0.44 % | 0.38 % |
| 0 km/h | 0.00 % | 0.00 % | 0.00 % | 0.00 % |
| 5 km/h | −0.53 % | −0.38 % | −0.29 % | −0.23 % |
| 10 km/h | −0.90 % | −0.60 % | −0.42 % | −0.30 % |
| 15 km/h | −1.13 % | −0.68 % | −0.46 % | −0.38 % |
| 20 km/h | −1.20 % | −0.75 % | −0.63 % | −0.60 % |
| 30 km/h | −1.50 % | −1.35 % | −1.41 % | −1.50 % |
| 40 km/h | −2.41 % | −2.56 % | −2.79 % | −3.00 % |

**Figure 1:**
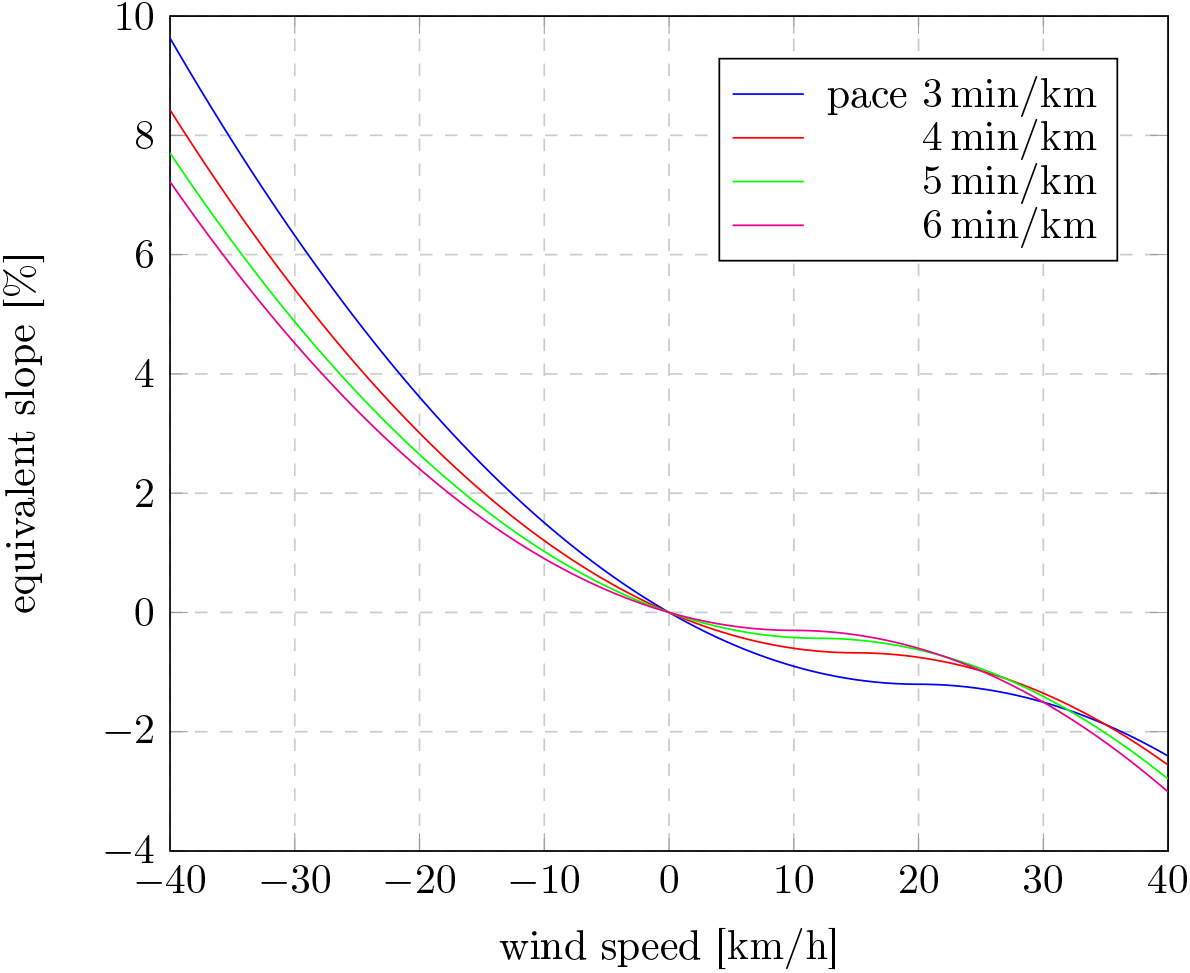
Equivalent slope in headwind and tailwind

Although two completely different forces are being compared in this consideration, the results are quite realistic. This can be explained by the fact that from a purely mechanical point of view runners have to deal with both forces, wind and gravity, in a very similar way. In the first case, the point of application of the force is at the geometric center of the cross section, while in the second case, it is located at the center of mass. Thus, in both cases, the force is applying at a similar body height.

The runner in the headwind will slightly increase his forward lean, and will therefore take a similar angle relative to the ground than the runner on an incline. During the flight phase, the runner is being pulled back in both cases in opposite running direction, i.e., tangential to the ground surface, and the corresponding losses must be compensated with every step by muscular work.

From a mechanical point of view, both situations are thus quite comparable. In practice though, additional effects may play a role. Headwind, for instance, can represent a psychological burden, since in some way, one is dealing with an invisible opponent. On the other hand, at high temperature, headwind can lead to cooling effects, which are not provided by a corresponding incline.

The mechanical comparability described here, is valid to the same extent for the considerations in section 2. The deceleration during the flight phase is being simulated on the treadmill by the fact that during the ground contact, the runner is loosing altitude, which he has to compensate for by muscular work. Non-mechanical aspects, like the absence of the cooling wind, but also the unnaturally high constancy of the speed, as well as the general monotony of running on the treadmill, will have more influence than possible mechanical differences.

## 4 Equivalent slope on a turning point route

In this section, we determine the average equivalent slope for a run at wind speed *w* on a turning point route, i.e., for covering the same track in both directions. In what follows, we will assume *w* ≥ 0 (tailwind) for the first half of the run, while for the second half, i.e., the way back, the speed will be assumed to be −*w* (headwind).

The additional energetical effort is given by the sum of the additional efforts of both sections, and the latters are proportional to the corresponding equivalent elevation gain. As a result, we have for the equivalent elevation gain of the complete track

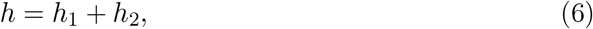

where *h*_*i*_ (*i* = 1, 2) denotes the elevation gain on the first and second half respectively. Denoting by *s*_*i*_ (*i* = 1, 2) the corresponding equivalent slopes, we obviously have *h*_*i*_ = *Ls*_*i*_*/*100, with *L* the length of the one way track. For the net slope *s* and the total elevation gain *h*, we have instead the relation *h* = 2*Ls/*100, because the length of the total track (round trip) is 2*L*. Comparing *h*_1_ + *h*_2_ to *h*, we find the equivalent slope for the total track in terms of the average value of the equivalent slopes of both halves, i.e.,

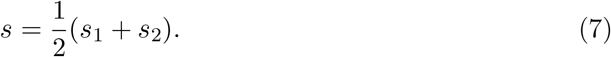

For the first half, we calculate *s*_1_ with the help of equation Eq. (5), while for the way back (*s*_2_), we use Eq. (5) with *w* → − *w*. For the average value, we get the following result:

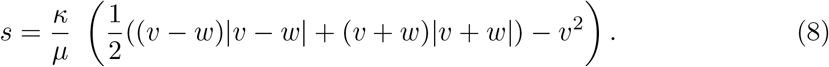

Considering the two possible signs of *v* − *w* (*v* + *w* is positive anyway, because *w* ≥ 0), we find the simpler representation

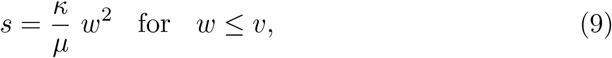

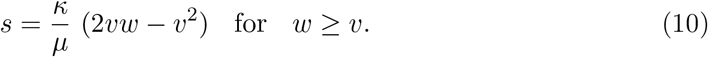

It’s not hard to convince oneself that *s* is always positive. The presence of wind on a turning point route always leads to an increased energetical effort.

In Table 3 and in Fig. 2, we display the results for paces between 3 min*/*km and 6 min*/*km, using the values for *κ* and *µ* given in section 1.

**Table 3:** Equivalent slope *s* on turning point route

| $w$ | 3:00 min/km | 4:00 min/km | 5:00 min/km | 6:00 min/km |
| --- | --- | --- | --- | --- |
| 0 km/h | 0.00 % | 0.00 % | 0.00 % | 0.00 % |
| 5 km/h | 0.08 % | 0.08 % | 0.08 % | 0.08 % |
| 10 km/h | 0.30 % | 0.30 % | 0.30 % | 0.30 % |
| 15 km/h | 0.68 % | 0.68 % | 0.65 % | 0.60 % |
| 20 km/h | 1.20 % | 1.13 % | 1.01 % | 0.90 % |
| 30 km/h | 2.41 % | 2.03 % | 1.73 % | 1.50 % |
| 40 km/h | 3.61 % | 2.93 % | 2.46 % | 2.11 % |

**Figure 2:**
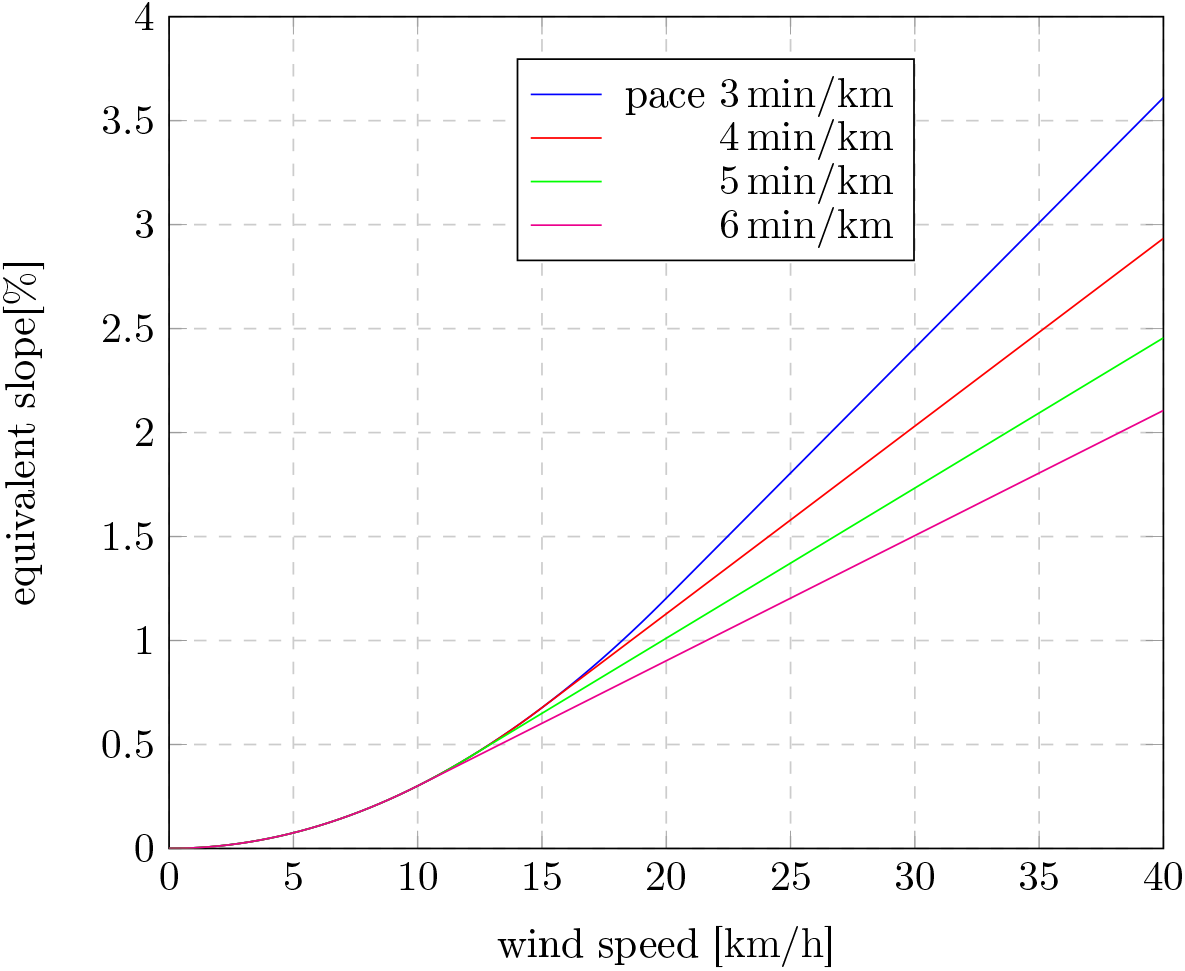
Equivalent slope on turning point route

It is interesting to note that the expected quadratic dependence of the wind speed only holds for wind speeds *w* smaller than the running speed *v*, see Eq. (9). For *w* ≥ *v*, we are dealing with a linear dependence, see Eq. (10).

The statement of the results of this section consists in showing that running on a flat turning point route with speed *v*, or with a corresponding pace, in presence of a wind speed *w* parallel to the running direction, is as *demanding* than running in the absence of wind, but facing instead an incline *s* during the complete run, i.e., on the way there and back. In both situations, comparable achievements should thus be obtainable. The results are based on purely mechanical considerations and will be refined in the next section.

## 5 Physiological aspects for the turning point route

The results obtained in the previous section are based on the consideration of purely mechanical aspects, while physiological aspects have been excluded entirely. This approach is justified for the situations described in sections 2 and 3, because for the moderate slopes or wind speeds under consideration, the physical challenges are to a great extent identical, as we have outlined at the end of section 3.

The physiological efforts, however, are not directly proportional to the mechanical ones. In particular, a negative slope doesn’t necessarily compensate for a positive slope of the same percentage, even though from an energetical point of view, this should be the case. Everybody knows from personal experience that better results can be achieved on flat tracks than on turning point routes with elevations, even though the net elevation cancels out.

For the calculations of section 4, this means that a positive slope *s* has to be rated higher than a negative slope of the same absolute value in order to obtain a physiologically meaningful *average* slope.

The correct weighting is hard to determine scientifically, and certainly depending on various physical and personal factors. However, good empirical approaches can be found in literature. We will now compare three alternatives for a concrete situation, two being based on online calculators, and the third one consisting of a *rule of thumb*.

From the results of section 4, we consider the case with pace 4 min*/*km at wind speed 15 km*/*h. Let the route consist of 5 km in one way (headwind) and 5 km back (tailwind). In section 3, we established an equivalent slope of *s* = 2.03 % for the headwind part and of − 0.68 % for the tailwind part (see Table 2). For the turning point route, we found a net equivalent slope of 0.68 % (see Table 3). Over the complete run, this leads to approximately 102 positive elevation meters, 34 negative elevation meters and 68 net elevation meters.

First, we will use the online calculator of Peter Greif, see [3]. If to a runner it takes 40 minutes on a flat track, then, according to the calculator, it will take him 42:10 for a track with 102 positive elevation meters and 34 negative ones. To the same runner, it will take 41:37 for a track with 68 positive elevation meters (and 0 negative ones). With the online calculator of lauftipps.ch [4], we find very similar results, namely 42:04 and 41:34 accordingly. Those times are supposed to represent an estimate on how long it will take the runner for the corresponding routes, without changing its physiological effort compared to the flat route.

A third method of calculation is based on the following rule of thumb: 100 positive elevation meters are equivalent to about 600 m additional distance, while 100 negative elevation meters correspond to a reduction of the distance of about 300 m.^3^ Running at a pace of 4 min*/*km, this corresponds to the addition of 144 s for 100 m uphill, and 72 s reduction for 100 m downhill. Applying this rule to our example, we find a running time of 42:02 for the route with 102 positive and 34 negative elevation meters, and 41:38 for the route with 68 elevation meters.

Again, we find very similar results, so for the rest of this section, we will stick to the rule of thumb, especially because the underlying formulas of the online calculators are not known. To be somewhat more general, we will, for the moment, denote the 600 m additional route for 100 elevation meters by *a*_2_, and the 300 m reduction of the distance per 100 negative elevation meters by *a*_1_.

In section 4, we calculated the equivalent slope for a turning point route at running speed *v* by considering the average value of the equivalent slopes of both halves of the route, see Eq. (7), with *s*_1_ ≤ 0 and *s*_2_ ≥ 0, thus assuming tailwind during the first half. We will now improve this average value in order to take into account the physiological differences present during both halves of the run.

According to the rule of thumb, we rescale the first half (negative elevation) with *a*_1_ and the way back (positive elevation) with *a*_2_ and write for the physiologically adjusted equivalent slope instead of Eq. (7) the following, weighted average

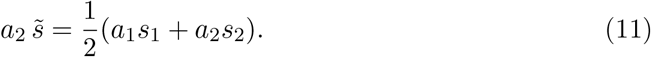

Since 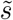 is always positive, we also have to rescale the left hand side of the equation with the factor *a*_2_. The final result is obtained after division by *a*_2_.

Let’s go back to our example of the 10 km route at a pace of 4 min*/*km and wind speed 15 km*/*h. From Table 2 in section 3, we take the values *s*_1_ = − 0.68 % and *s*_2_ = 2.04 % for both parts of the track. Using *a*_1_ = 300 and *a*_2_ = 600, we find 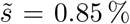 for the adjusted equivalent slope of the complete track, which is higher then the unadjusted value *s* = 0.68 % from Table 3.

To check the calculations, we can apply the rule of thumb once again to the above example. For the track with 85 elevation meters (resulting from 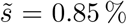 for 10 km), we now get 0.85 · 600 additional meters, corresponding to 122 additional seconds and thus to an adjusted running time of 42:02. This corresponds exactly to the result we found by applying the rule of thumb to the track with 102 positive and 34 negative elevation meters.^4^

In this sense, for a runner running at a pace of 4 min*/*km, a route with a slope of 0.85 % is physiologically equivalent to a route with a slope of 2.04 % on one half and a slope of − 0.68 % on the other half, and thus equivalent to a flat turning point route at a wind speed of 15 km*/*h.

Inserting the slopes of both parts explicitly into Eq. (11), we get

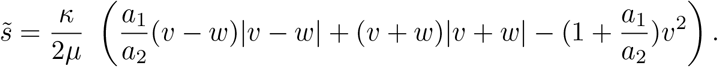

Considering again the two possible signs of *v* − *w*, we find the slightly simpler representation

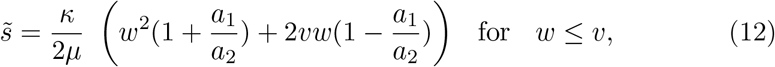

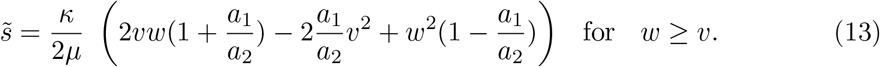

In the case *a*_1_*/a*_2_ = 1, this reduces to Eqs. (9) and (10). However, as we have seen, *a*_1_*/a*_2_ = 300*/*600 = 1*/*2 corresponds to a physiologically more realistic value.

Instead of Table 3, we get from Eqs. (12) and (13) the physiologically adjusted Table 4. The plot in Fig. 3 shows the adjusted results for various running speeds. The difference between the adjusted and the unadjusted equivalent slopes is shown in Fig. 4.

**Table 4:** Physiologically adjusted equivalent slope 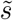 on turning point route

| $w$ | 3:00 min/km | 4:00 min/km | 5:00 min/km | 6:00 min/km |
| --- | --- | --- | --- | --- |
| 0 km/h | 0.00 % | 0.00 % | 0.00 % | 0.00 % |
| 5 km/h | 0.21 % | 0.17 % | 0.15 % | 0.13 % |
| 10 km/h | 0.53 % | 0.45 % | 0.41 % | 0.38 % |
| 15 km/h | 0.96 % | 0.85 % | 0.77 % | 0.70 % |
| 20 km/h | 1.51 % | 1.32 % | 1.17 % | 1.05 % |
| 30 km/h | 2.78 % | 2.37 % | 2.09 % | 1.88 % |
| 40 km/h | 4.21 % | 3.57 % | 3.15 % | 2.86 % |

**Figure 3:**
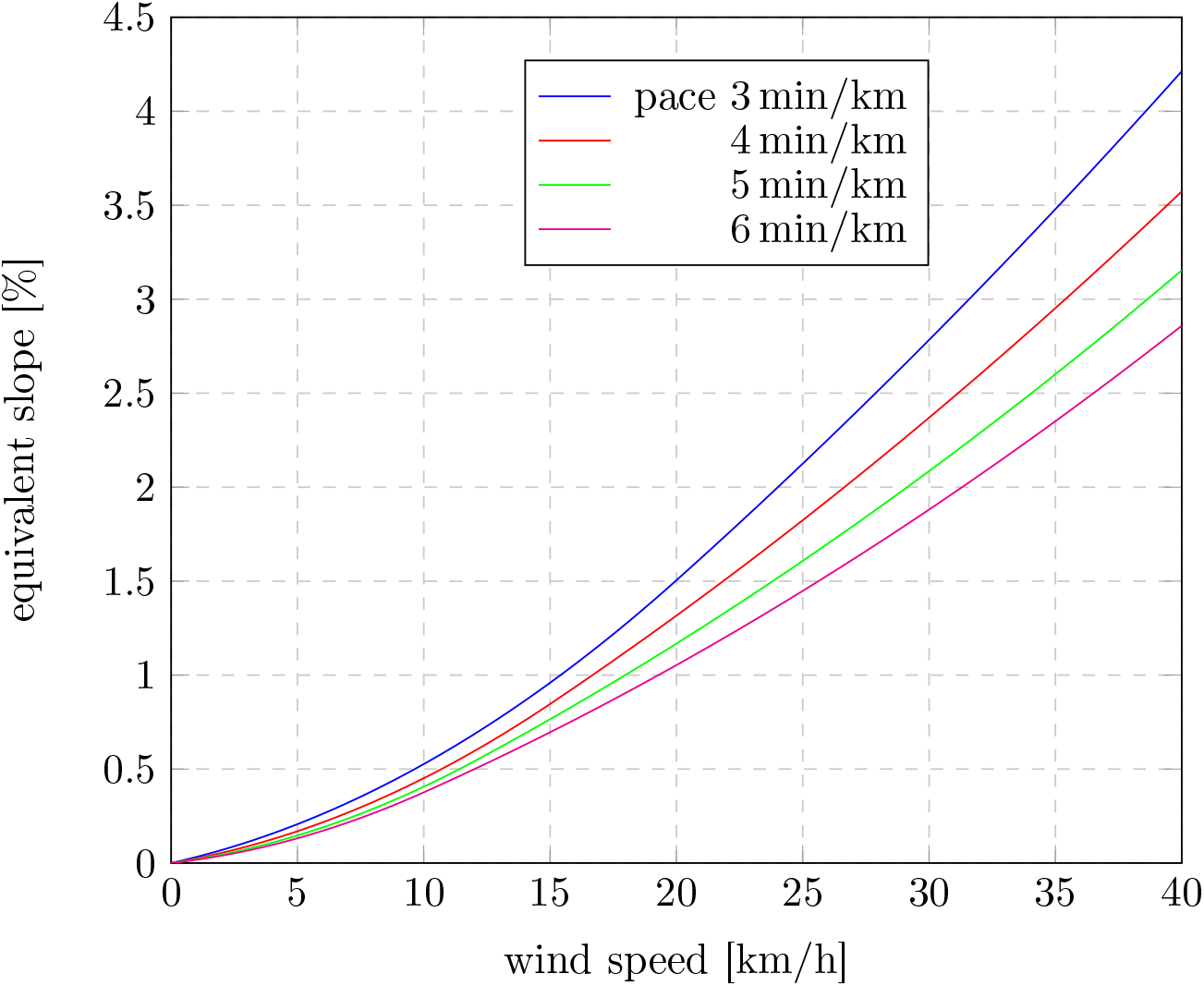
Adjusted equivalent slope on turning point route

**Figure 4:**
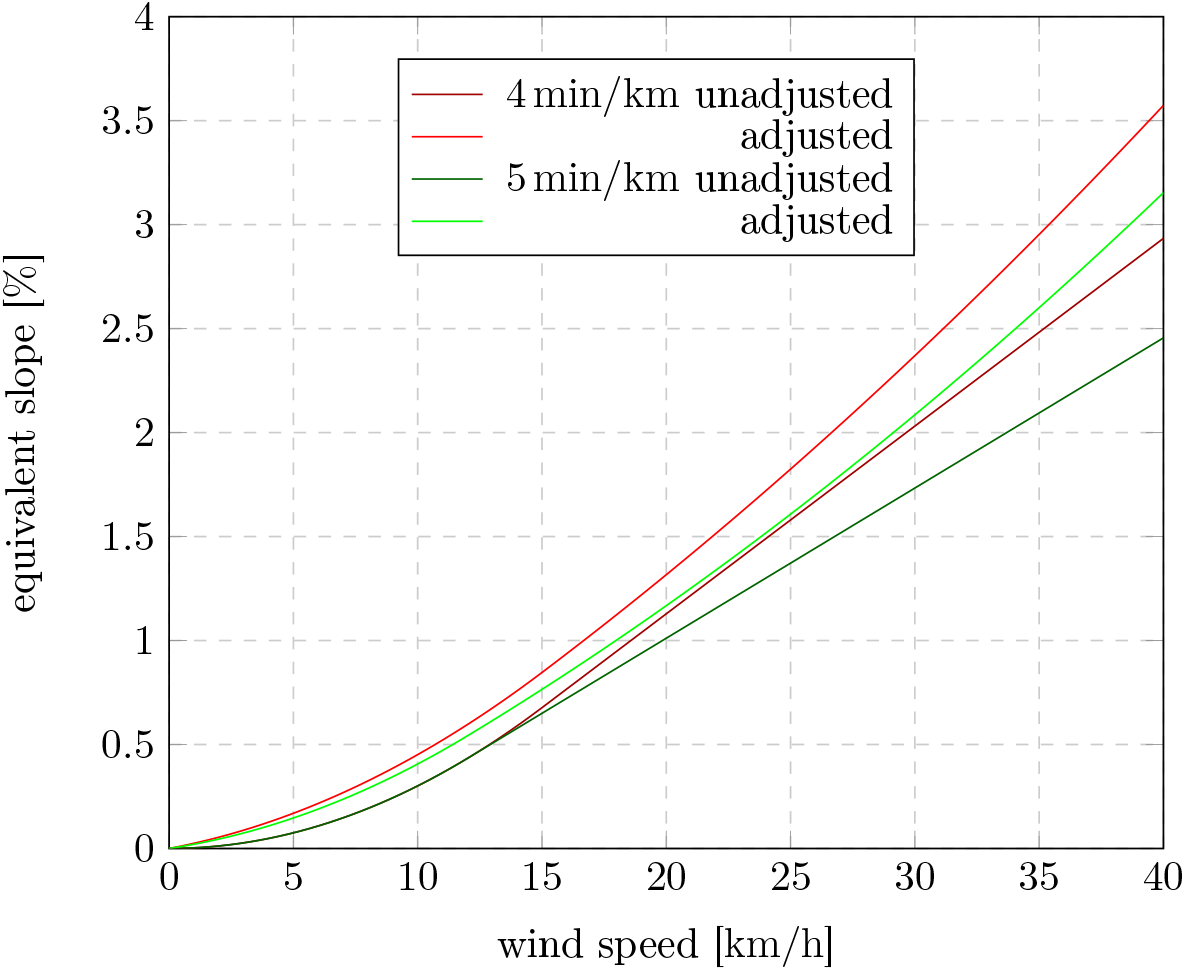
Unadjusted vs. adjusted equivalent slope

Apparently, the effects are clearly noticeable already at low wind speeds.

## 6 Adjustment of the running speed

In view of the nonlinear velocity dependence of the air resistance force, it seems obvious that by an appropriate adjustment of the running speed, the energetical effort on a turning point route can be reduced. The idea consists in running the tailwind part faster than the headwind part, in order to reduce the relative velocities that enter the force, and thus the total energy consumption (force times distance), quadratically.

In this section, we investigate whether this is indeed possible, and which are the optimal speeds for both directions.

As before, we assume tailwind for the first half of the route. Let *v*_1_ and *v*_2_ be the running speeds of the first and second half respectively, and *v* the average speed for the complete run. We thus have for the corresponding time intervals:

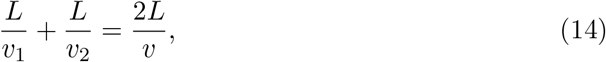

where *L* denotes the length of the one-way track. From Eq. (1), the force, and thus the energy effort on the given distance, is proportional to 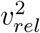:

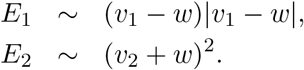

The task consists in finding *v*_1_ and *v*_2_ minimizing the sum *E*_1_ + *E*_2_. To this end, we insert *v*_2_ from Eq. (14), i.e., *v*_2_ = *vv*_1_*/*(2*v*_1_ − *v*), into the expression for *E*_1_ + *E*_2_, take the derivative with respect to *v*_1_ and determine its root. Since we are dealing with an equation of fourth degree, the root can be written down in closed form. The corresponding expression, however, is very complicated. For this reason, we confine ourselves to a representation of the numeric results in form of a table, see Table 5. For convenience, the velocities have been transformed to paces (min*/*km) and refer to the tailwind part.^5^

**Table 5:** Optimal pace for tailwind part

| $w$ | 3:00 min/km | 4:00 min/km | 5:00 min/km | 6:00 min/km |
| --- | --- | --- | --- | --- |
| 0 km/h | 3:00 min/km | 4:00 min/km | 5:00 min/km | 6:00 min/km |
| 5 km/h | 2:45 min/km | 3:33 min/km | 4:19 min/km | 5:01 min/km |
| 10 km/h | 2:30 min/km | 3:08 min/km | 3:39 min/km | 4:05 min/km |
| 15 km/h | 2:16 min/km | 2:43 min/km | 3:04 min/km | 3:18 min/km |
| 20 km/h | 2:03 min/km | 2:22 min/km | 2:34 min/km | 2:42 min/km |

Before we discuss the results, we will take a look at the energy saving obtained by those speed adjustments. Similar to the considerations in section 4, an equivalent scope can be calculated for the adjusted velocities. To this end, we split Eq. (8) into the tailwind and headwind parts and insert the corresponding velocities. We obtain the following equations:

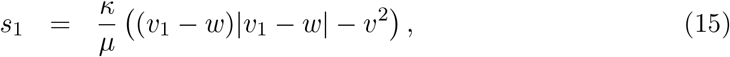

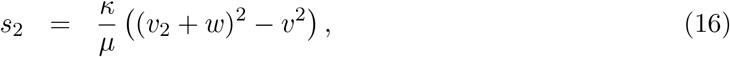

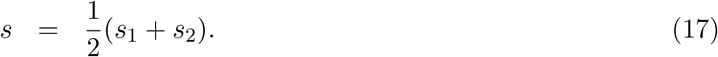

The terms proportional to − *v*^2^ take into account that we are interested in the additional effort, compared to the windless route at constant speed *v*, just as in Eq. (8). The results are displayed in Table 6. For convenience, the equivalent slopes corresponding to the unadjusted (constant) speed *v* from Table 3 are indicated in brackets.

**Table 6:** Equivalent slope for turning point route at optimal pace

| $w$ | 3:00 min/km | | 4:00 min/km | | 5:00 min/km | | 6:00 min/km | |
| --- | --- | --- | --- | --- | --- | --- | --- | --- |
| 0 km/h | 0.00 % | (0.00 %) | 0.00 % | (0.00 %) | 0.00 % | (0.00 %) | 0.00 % | (0.00 %) |
| 5 km/h | 0.05 % | (0.08 %) | 0.05 % | (0.08 %) | 0.05 % | (0.08 %) | 0.05 % | (0.08 %) |
| 10 km/h | 0.20 % | (0.30 %) | 0.20 % | (0.30 %) | 0.20 % | (0.30 %) | 0.20 % | (0.30 %) |
| 15 km/h | 0.47 % | (0.68 %) | 0.44 % | (0.68 %) | 0.44 % | (0.65 %) | 0.44 % | (0.60 %) |
| 20 km/h | 0.79 % | (1.22 %) | 0.78 % | (1.14 %) | 0.77 % | (1.02 %) | 0.76 % | (0.91 %) |

We see that by an optimal choice of the running speed for both halves of the turning point route, a considerable reduction of the equivalent slope, and thus of the energetical effort, can indeed be achieved. On the other hand, Table 5 also shows that the calculated velocities for the tailwind part are extremely high and in most cases unrealistic.

To obtain realistic values for the optimal velocities, one has to take into account additional effects. Our calculations are based merely on air resistance and gravity. The basic effort of running as it occurs, e.g., on a flat treadmill, has not been taken into account, or more accurately, has been assumed to be speed independent. This effort is basically a result of the permanent accelerations and changes of direction of the runner’s arms and legs, as well as of the vertical up and down movements of the body, with energy losses during the impact of the foot, see, e.g., [5].

This effort is increasing with speed, not only per time unit (higher power), but also in terms of a higher total energy consumption. For a 10 km run on a treadmill in 40:00, more energy is needed than for the same run in 50:00. Moreover, the dependence of this effect from the velocity is nonlinear, meaning that slower segments are not energetically compensated by faster ones.^6^

In addition, physiological effects can lead to a decrease in efficiency and an increase of fatigue effects with increasing performance (power). For this reason, e.g., a runner will decrease his speed when running uphill, and increase it when running downhill. From the point of view of physics, it would be preferable to run at constant speed, not only in view of air resistance effects, but also because of the basic effort just described.

In general, these effects are opposing each other. The overall energetical optimum (including the basic effort) on a turning point route, be it in the mountain or in the wind, can mathematically lead to a speed distribution which lies partly in an unfavorable, too high performance range. The physiological optimum, i.e., the optimum compromise between minimum energy consumption and a balanced power output, depends on personal aspects, for example on the efficiency curve of the runner in terms of performance, but also on the length of the route and the endurance of the runner, and ultimately remains a matter of experience.^7^

Quite generally, the runner will choose an intermediate way: He will increase the effort on the uphill part, but not to the extent that he maintains the same speed than on a flat track. Similarly, he will reduce power running downhill, but will still run faster than on a flat track.^8^

In the same way, the runner on a turning point route should decrease his speed on the headwind part and increase it on the tailwind part. Besides the optimization of the power balance, in the case of air resistance, this provides a direct energetical advantage, as we have outlined at the beginning of this section. However, as we have argued, the theoretically optimal speed adjustments are not realistic, as they lead to extreme velocities.

To get an impression of the realistically possible optimization, we assume that the runner decreases (increases) his pace by one second per km*/*h wind speed in headwind (tailwind). Thus, e.g., at a wind speed of 10 km*/*h, a runner who runs at a pace of 4 min*/*km in the absence of wind, will be running at a pace of 3:50 min*/*km in tailwind and of 4:10 min*/*km in headwind. Such speed adjustments can be considered to be realistic and correspond to some extent with everyday experience.

In Table 7, the equivalent slopes for such speed adjustments, calculated according to Eqs. (15)-(17), have been summarized. As expected, the values are in between those obtained without pace adjustment, see Table 3 in section 4, and those that would be obtained for the energetically optimal (as far as the air resistance is concerned) pace adjustment, see Table 6.

**Table 7:** Equivalent slope for turning point route with realistic pace adjustment

| $w$ | Δ-pace | 3:00 min/km | 4:00 min/km | 5:00 min/km | 6:00 min/km |
| --- | --- | --- | --- | --- | --- |
| 0 km/h | $\pm 0$ s | 0.00 % | 0.00 % | 0.00 % | 0.00 % |
| 5 km/h | $\pm 5$ s | 0.06 % | 0.07 % | 0.07 % | 0.07 % |
| 10 km/h | $\pm 10$ s | 0.24 % | 0.27 % | 0.28 % | 0.28 % |
| 15 km/h | $\pm 15$ s | 0.55 % | 0.60 % | 0.61 % | 0.58 % |
| 20 km/h | $\pm 20$ s | 0.98 % | 1.03 % | 0.96 % | 0.87 % |

We will not present physiologically adjusted values, like the ones in section 5, Table 4, because, as has been outlined above, the speed adjustment itself gives rise to additional physiological effects that are not covered by our rule of thumb.

Roughly, for the adopted speed adjustment, the energy optimization, and thus the reduction of the equivalent slope amounts to about 20 % for a pace of 3 min*/*km, 10 % for 4 min*/*km, 6 − 7 % for 5 min*/*km and only 3 − 6 % for 6 min*/*km.^9^ For wind speeds below 10 km*/*h, the optimization rate is even higher.

In practice, this means that by choosing such a speed adjustment on a turning point route, the influence of the wind can be reduced by the corresponding rate. Physiological aspects, like those considered in section 5, or additional effects arising as a result of the speed adjustment, have been neglected.

It should be noted that the choice of the adjustment merely serves as an example and does not necessarily represent the physiologically optimal adjustment, which cannot be determined in general. Contrary to the theoretical results in Table 6, being based on purely mechanical arguments, the results of Table 7 are likely to provide a realistic insight on the effect of the speed adjustments that will ultimately be chosen *instinctively* by the runner.

## 7 Summary

The effects of headwind and tailwind have been compared with the effects of running uphill or downhill. To this end, the influence of the wind has been numerically expressed in terms of *equivalent slopes* for several situations:

- running on a treadmill,
- running in tailwind or headwind,
- running on a turning point route,
- running on a turning point route with optimized speed.

On a purely mechanical level, the additional forces arising while running up or down an incline have been compared with those arising in the presence of headwind or tailwind.

As long as the run is being performed in one direction (no changes in the wind speed), and as long as the incline and the running speed are constant throughout the complete run, all additional physiological aspects are, in sufficient approximation, comparable. In those cases, the mechanically arising additional forces, i.e., the gravitational force on one hand and the increased or decreased air resistance on the other hand, can be directly used to determine the corresponding equivalent slopes. The related calculations have been performed for running on a treadmill (section 2) as well as for running in tailwind or headwind (section 3). The corresponding results are summarized in Table 1 and Table 2, as well as in the plot in Fig. 1.

In section 4, the same calculations have been applied to the turning point route, see Table 3 and Fig. 2 for the results. Due to physiological effects, however, the equivalent slope for the complete track cannot be assumed to be equal to the arithmetic mean of the equivalent slopes of both halves of the track. This aspect has been taken into account in section 5. In Table 4 and in Fig. 3, physiologically adjusted results for the turning point route have been presented.

Finally, in section 6, the influence of speed adjustments on the energy balance for the turning point route has been investigated. Keeping the average speed (and thus, the total running time) constant, possible optimizations achieved by choosing different speeds for both halves of the track have been estimated. As far as the air resistance is concerned, energetically optimized paces for both parts can be computed. However, those purely mechanical considerations lead to physiologically nonsensible, or even non realizable paces, see Table 5. For this reason, the energy optimizations, in form of the corresponding equivalent slopes, have been calculated assuming realistic speed adjustments instead. The results are summarized in Table 7.

## Footnotes

1 It should be noted that the slope indication of most treadmills is erroneous (if no calibration has been performed), and it is preferable to determine the slope by external means, e.g., with a spirit level with inclinometer.

2 We talk about tailwind whenever *w >* 0, even if *v* ≥ *w*, i.e., even if the ambient wind does not completely compensate for the *wind* produced by the runner himself.

3 We don’t know the exact origin of this rule, it can be found on many websites. Instead of 600 additional meters, some authors take 500 m, while for the corresponding reduction, values between 200 m and 400 m are being used.

4 Similar results can be obtained by applying the online calculators to a 10 km track with 85 elevation meters.

5 The correspondent paces for the headwind part can be deduced from the fact that the average pace is given by the arithmetic mean of the paces for both ways, which is, e.g., not the case for the corresponding velocities, see Eq. (14). Thus, for instance, if the turning point route has to be done at an average pace of 5 min*/*km, and for the tailwind part, we find 4:19 min*/*km in the table, then we will need a pace of 5:41 min*/*km for the headwind part.

6 In a concise formulation, this means, e.g., that an interval training is more demanding than a constant speed run over the same distance in the same time, even when wind effects can be neglected.

7 The rule of thumb used in section 5 represents an example of this situation. As a result of the physiological aspects, the runner will always suffer more losses running uphill than he will gain back on the downhill part, i.e., we always have *a*_2_ ≥ *a*_1_.

8 The power adjustments can easily be tracked with a heart rate monitor, albeit with a certain temporary delay.

9 It should be noted that the pace adjustments adopted here (1 s per km*/*h wind speed) are, in relative terms, higher for faster paces. For slower paces, stronger adjustments could be considered, leading to higher optimization rates.

